# piRNAs of *Caenorhabditis elegans* broadly silence nonself sequences through functionally random targeting

**DOI:** 10.1101/2021.07.31.454584

**Authors:** John McEnany, Yigal Meir, Ned Wingreen

## Abstract

Small non-coding RNAs such as piRNAs serve as guides for Argonaute proteins, enabling sequence-specific, post-transcriptional regulation of gene expression. The piRNAs of *Caenorhabditis elegans* have been observed to bind targets with high mismatch tolerance, and appear to lack specific transposon targets, unlike piRNAs in *Drosophila melanogaster* and other organisms. These observations support a model in which *C. elegans* piRNAs provide a broad, indiscriminate net of silencing, acting in competition with siRNAs associated with the CSR-1 Argonaute that specifically protect self genes from silencing. However, the breadth of piRNA targeting has not been subject to in-depth quantitative analysis, nor has it been explained how piRNAs are distributed across sequence space to achieve complete coverage. Through a bioinformatic analysis of piRNA sequences, incorporating an original metric of piRNA-target distance, we demonstrate that *C. elegans* piRNAs are functionally random, in that their coverage of sequence space is comparable to that of random sequences. By possessing a sufficient number of distinct, essentially random piRNAs, *C. elegans* is able to target arbitrary nonself sequences with high probability. This result elucidates the mechanism by which newly transcribed mRNAs in *C. elegans* are classified as self or nonself, and has implications for piRNA evolution and biogenesis.

## INTRODUCTION

Piwi-interacting RNAs, or piRNAs, are small RNAs often implicated in silencing of transgenes and deleterious transposons. The characteristics of piRNA targeting behavior differ among organisms (for a review, see Parhad and Theurkauf (1)). In *D. melanogaster*, piRNA sequences serve as a library of transposon subsequences, which enables silencing through sequence-specific target identification (2, 3). In the nematode *C. elegans*, however, most piRNAs appear to lack complementarity to transposons — or indeed, any other clear target in the genome. Instead, researchers propose that the spectrum of *C. elegans* piRNAs can target virtually *any* reasonably-sized mRNA sequence (4,5). Evidently, such broad silencing must be coupled with a licensing system to maintain a proper transcription profile. What can a quantitative understanding of piRNA targeting reveal about the functionality and evolution of this system, combining broad silencing with specific licensing? How do piRNAs achieve such broad sequence coverage, and is their capacity to target nonself sequences at all dependent on the piRNA sequences themselves?

The piRNAs of *C. elegans* are 21 nucleotides in length and serve as guide RNAs for the PRG-1 Argonaute, which triggers the RNA interference pathway to initiate epigenetic silencing of its targets (6,7). In *prg-1* knockouts, sterility occurs in a temperature-dependent manner (8) or across the course of multiple generations (9), illustrating the importance of the piRNA system in maintaining proper gene expression. The expression of self-genes must be protected from piRNA-mediated silencing via a licensing system, which is not yet fully understood. One promising candidate for a licensing Argonaute is CSR-1, which binds siRNAs complementary to most germline-expressed genes, and when recruited to a transcript, protects it from piRNA-mediated silencing (10,11). A schematic of the putative *C. elegans* self/nonself discrimination system is presented in Figure 1. The licensing system maintains a memory of self-genes, allowing the piRNA-mediated silencing system to broadly target nonself sequences. This broad targeting is largely due to the mismatch-tolerant pairing rules between piRNAs and their targets. Studies of piRNA-mRNA crosslinking, and of the targets of synthetic piRNAs, have identified a “seed region” at nucleotides 2-8 which is particularly important for targeting, along with a potential supplementary region at nt 14-19 (5,4) (Fig. 1). Outside of these regions, piRNAs tolerate significant amounts of non-canonical base pairing — particularly GU wobbles, which appear less heavily penalized than other mismatches (5).

**Figure 1.**
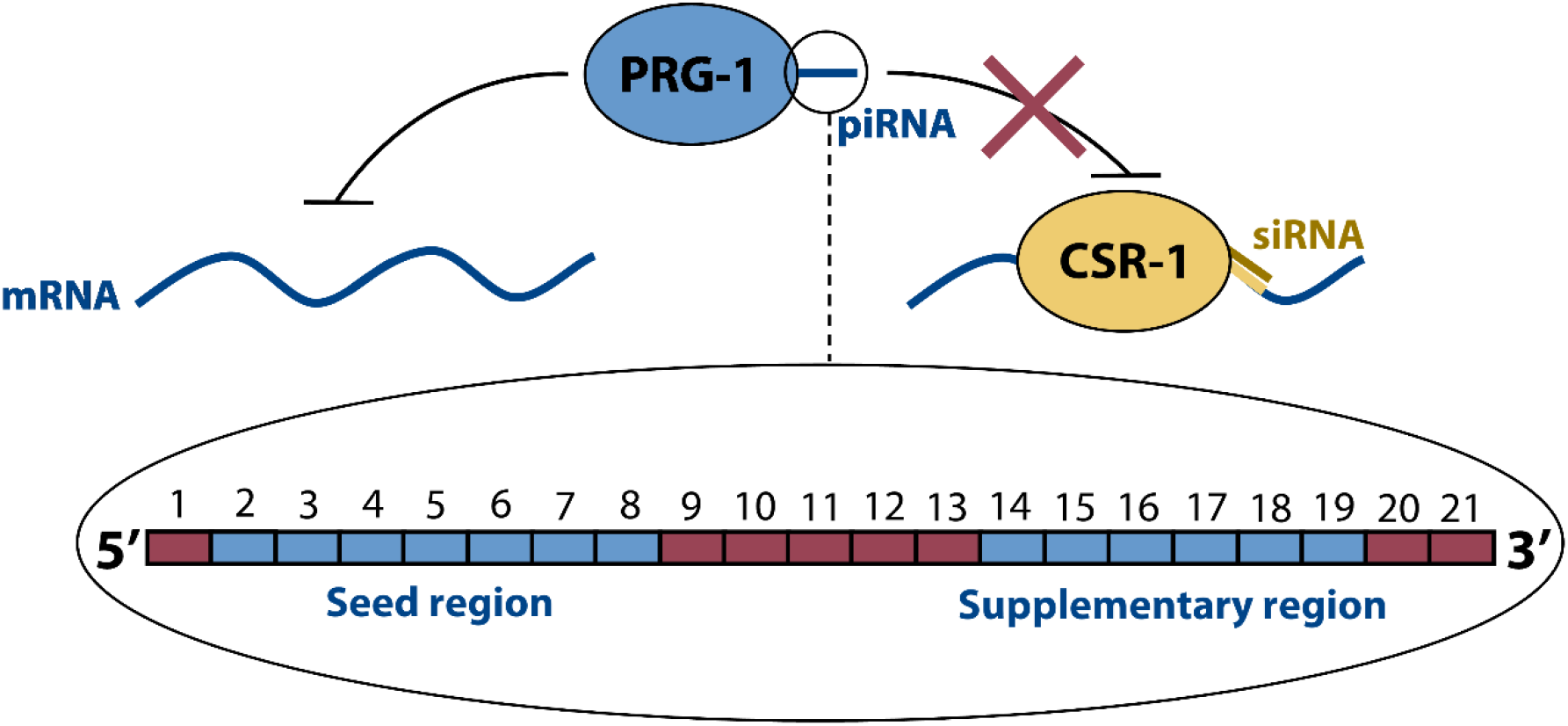
Schematic of the putative *C. elegans* self-nonself discrimination system, as mediated by small RNAs. PRG-1 Argonautes are guided by piRNAs to matching target sequences on self and nonself mRNA transcripts. As shown in the enlarged view, each piRNA is 21 nucleotides long, with a seed region at nt 2-8 and a supplementary region at nt 14-19 where canonical base pairing is particularly important for target recognition. When a PRG-1:piRNA binds to a valid target, RNA interference and downstream silencing are initiated. CSR-1 Argonautes are guided by siRNAs to matching sequences on self-transcripts. Binding by CSR-1:siRNA licenses a transcript, protecting it from silencing by PRG-1.

The theory that *C. elegans* piRNAs engage in broad, non-specific targeting is relatively well-supported by data. A cross-linking, ligation and sequencing of hybrids (CLASH) study revealed that piRNAs interact with a wide variety of germline mRNAs, with single piRNAs having numerous targets and single mRNAs possessing multiple potential piRNA binding sites (4). A bioinformatic analysis which used a formalization of piRNA pairing rules to identify possible targeting sites found that about half of germline-expressed genes had a piRNA targeting site, and calculated similar proportions for germline-silenced genes and control sequences (5). However, the mechanism through which piRNAs achieve their coverage of sequence space is still unclear: their sequences could minimize redundancy (maximizing their coverage with the minimum number of distinct sequences), or simply be effectively random, but exist in large enough numbers that at least one piRNA will have a suitable site on any target mRNA. Furthermore, it is unclear to what extent piRNA sequences match transposon sequences or avoid sequences of self-genes; despite the lack of clear matches, it is possible that piRNA sequences are more likely to target transposons than self-transcripts when considering the permissive piRNA targeting rules. Bagijn et al. (7), for instance, found that putative piRNA target loci are depleted of protein-coding genes but not transposons. Answering these questions is vital to developing a more complete understanding of the *C. elegans* genome defense system. In particular, the likelihood of a given nonself sequence being targeted by a piRNA can be used to investigate the timescale and accuracy of self/nonself discrimination.

To address these questions, we conducted a bioinformatic analysis of *C. elegans* piRNAs, by developing a distance metric based on the log-odds likelihood of observing a sequence of matches, mismatches and GU wobbles in a piRNA-target pair. We found that piRNA sequences are not self-avoiding (i.e. optimized to avoid redundancy in sequences), and are instead essentially random — but present in a large enough number to cover all of sequence space with a high probability. Furthermore, *C. elegans* piRNAs are functionally equivalent with respect to transposon and self-transcript sequences, with only minor differences from random controls in terms of their targeting behavior. We propose that *C. elegans* piRNAs evolved to be essentially random, employing a strategy that enables targeting of a wide variety of nonself sequences.

## MATERIALS AND METHODS

### Quantification of sequence matches

To quantify the sequence matches between a set of piRNAs and a set of transposon or transcript sequences, every piRNA was checked against every same-length subsequence on each transposon or transcript, and the number of mismatches at the best possible binding site was recorded. The proportion of mismatched bases (i.e., non-canonical base pairs) was used to quantify dissimilarity for both *C. elegans* and *D. melanogaster*. Throughout our analysis, we compared real piRNAs to random control sequences. The latter were generated to be the same lengths (21 nt for *C. elegans*, 15-30 nt for *D. melanogaster*) and number (17,849 and 13,904, respectively) as the original sets of piRNAs, but with randomized sequences. These sequences were generated pseudo-randomly such that the probability of a given nucleotide was equal to the probability of the nucleotide appearing in a real piRNA from the organism. (*C. elegans*: 31% A, 36% U, 16% C, 17% G. *D. melanogaster:* 20% A, 24% U, 21% C, 35% G. For *C. elegans*, we omitted the first nucleotide, usually U, when calculating these probabilities.) We also measured sequence dissimilarity between *C. elegans* piRNAs and their potential mRNA targets using a functional piRNA distance metric, as detailed in the next section.

### piRNA-piRNA and piRNA-target distance

We developed a functional piRNA distance metric, taking into account the importance of the position of each base for piRNA targeting. This metric is based on a log-likelihood estimate of observing a match, mismatch or GU wobble at each base in experimentally confirmed piRNA-target pairs, meaning that bases which are more likely to match in known piRNA-target pairs were weighted more heavily when calculating the distance. Past research has suggested that GU wobbles are more favorable for piRNA targeting than other mismatches, so we constructed the metric such that a GU wobble could induce less of a distance penalty (but not more) than an arbitrary mismatch (5). We defined the piRNA-mRNA distance as

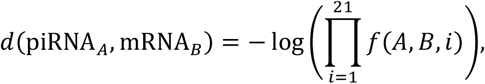

where

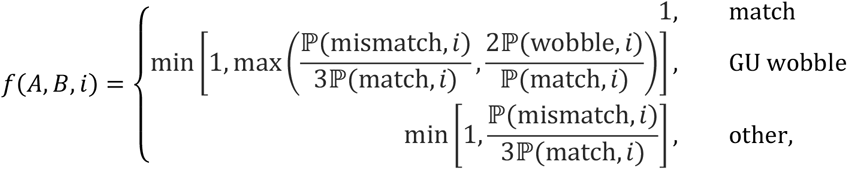

and the probabilities ℙ represent the probability of a canonical base pairing match, a GU wobble, or any non-canonical mismatch at each of the 21 sites. These probabilities were estimated from the 17 experimentally observed piRNA-target pairs used in Zhang et al., with a pseudocount of 1/16 for each of the 16 possible nucleotide pairs at each site (5). In order to verify that the number of experimentally confirmed pairs was sufficient to construct a self-consistent distance metric, we constructed leave-one-out test sets, calculating the piRNA-target distance for each of the 17 pairs while omitting that pair when calculating the probabilities. We compared these distances to those obtained from using all 17 piRNAs (the full training set), and took a small deviation to mean that the distance was self-consistent (Supplementary Figure 1). In addition to the piRNA-target distance, we used a similar metric to compare piRNAs with each other, based on the ability of each piRNA to target the other’s perfect complement. This piRNA-piRNA distance is:

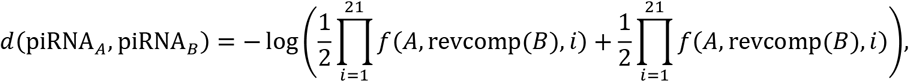

where revcomp denotes the reverse complement of a sequence. Using this metric, we calculated the distribution of *C. elegans* piRNA-piRNA distances using every pair of piRNAs. This analysis also allowed us to cluster piRNAs into groups of similar piRNAs, using hierarchical clustering with single linkage (in other words, defining the distance between two groups of piRNAs as the smallest distance between any pair of their members). A distance cutoff of 10 was used to define separate groups.

### Global piRNA targeting

In addition to analyzing pairs of transposons (or self-transcripts) and piRNAs, we also sought to investigate how well a transposon or transcript could be targeted by a set of piRNAs. Specifically, we used the techniques described to identify the smallest piRNA-target distance between the best-matching site on a transposon or transcript and the entire set of *C. elegans* piRNAs. To compare these results to a random control, we generated pairs of randomized 21-nt sequences representing “genes” and “piRNAs” (with an equal chance for each nucleotide), providing a random sample of piRNA-target distances. In order to adequately sample the left tail of the distribution of distances (corresponding to well-matching sequences), we oversampled the rare sequences with a high number of matches and then compensated by using the binomial probability of a random pair of sequences having a given number of matches. Specifically, *k* bases in each pair were randomly chosen to match, with *k* = 0…20, while the other bases were allowed to be GU wobbles or mismatches based on the corresponding probabilities expected for random sequences. For each value of *k*, we generated 10^7^ such randomized sequences. Then, we calculated the empirical cumulative distribution function (CDF) of distances for each set of 10^7^ sequences, which closely approximates the true CDF of the piRNA-target distance for random sequences conditioned on the number of matches. We then added these 20 CDFs, together with the *k* = 21 complete-match (zero-distance) case, each weighted by the binomial probability of two 21-nt sequences having *k* matches. This calculation produced the non-conditional CDF of the piRNA-target distance between two random 21-nt sequences:

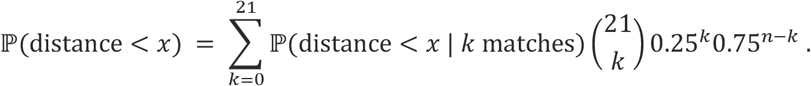

In order to determine the CDF of the shortest piRNA-target distance between a random “gene” of length *L* and a set of *N* random “piRNAs,” we assumed the different binding sites on a gene to be independent. This assumption allowed us to take the shortest piRNA-target distance as the minimum of *N* · *L* samples from the above single-pair distribution of distances. The distribution of this minimum can be calculated from the original CDF with the equation

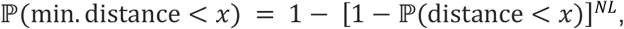

which allowed us to determine the predicted mean distance and the full probability density function for arbitrary choices of gene length and piRNA number. For ease of presentation, the probability density function obtained by numerically differentiating the cumulative distribution function was smoothed with a smoothing spline.

### Computational resources

*C. elegans* piRNA sequences were downloaded from piRTarBase at cosbi6.ee.ncku.edu.tw/piRTarBase/download/ (12). Full-length transcript sequences for *C. elegans* were obtained from WormBase ParaSite at parasite.wormbase.org/Caenorhabditis_elegans_prjna13758/Info/Index (13,14). *C. elegans* transposon sequences were obtained from Laricchia et al., Supplemental Table S16 (15). *D. melanogaster* piRNA sequences associated with the Piwi Argonaute were obtained from Brennecke et al., with GEO accession number GSM154620 (2). Transposon sequences for *D. melanogaster* were taken from the Natural Transposable Element Project (version 9.4.1) of the Berkeley *Drosophila* Genome Project, at fruitfly.org/p_disrupt/TE.html (16).

## RESULTS

### piRNA-transposon match comparison between *C. elegans* and *D. melanogaster*

*C. elegans* is thought to employ a strategy of transposon silencing which utilizes mismatch-tolerant piRNA binding to achieve broad targeting (4,5). By contrast, *D. melanogaster* relies on complementary piRNAs matching target transposon sequences (2,3). Before beginning an in-depth analysis of *C. elegans* piRNAs, we sought to quantitatively verify this key difference as a proof of concept. To use a metric consistent across organisms, we identified the best-matching site of each piRNA along each transposon for both organisms and counted the proportion of mismatches. We conducted the same analysis for real piRNAs and random controls, quantifying how biased piRNA sequences are toward targets on transposons in each organism (Figure 2).

**Figure 2.**
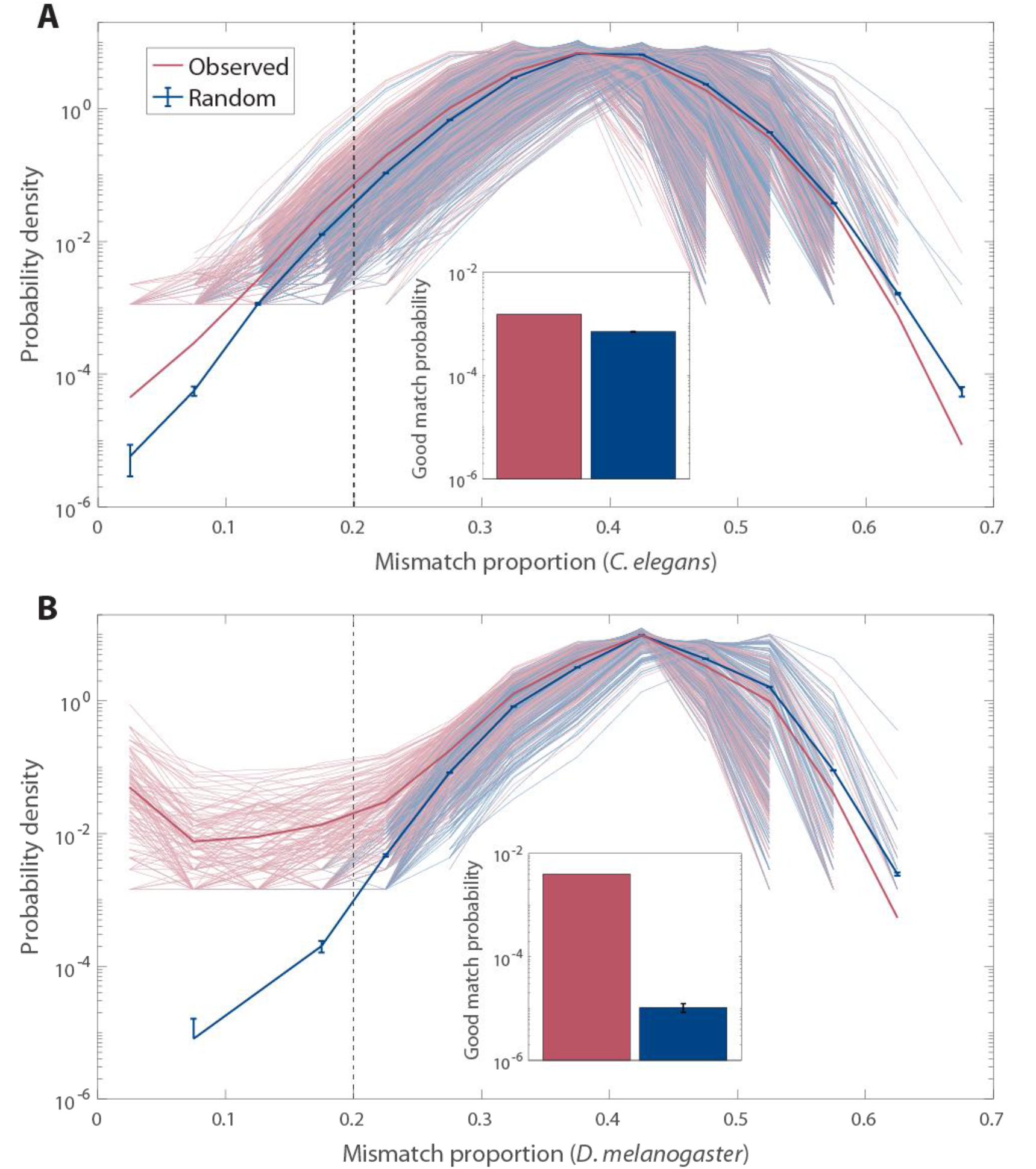
Bioinformatic comparison between *C. elegans* and *D. melanogaster* of piRNA-transposon sequence similarity. **(A)** *C. elegans* probability distribution of the proportion of mismatches within the best-matching site for every pair of piRNAs (*n* = 17,849) and transposons (*n* = 752); results shown for real piRNAs and random control sequences. Pale curves indicate the distribution for each individual transposon; solid curve is the average across all pairs. (For mismatch proportions that do not occur in the sample, no data is shown.) Inset: proportion of piRNA-transposon pairs where the best-matching site has fewer than 20% mismatches (left of the dashed gray line in the main panel), for real piRNAs and random control sequences. Error bars indicate counting error (square root of the number of counts). **(B)** Same as (A), but for *D. melanogaster* piRNAs (*n* = 13,904) and transposons (*n* = 179).

The difference between real and control piRNAs is much starker for *D. melanogaster* than *C. elegans*, particularly in the left tail (well-matching end) of the distribution of mismatches. While real *C. elegans* piRNAs are only about twice as likely as random controls to be “good” matches to a given transposon sequence (defined as matching any subsequence with fewer than 20% mismatches), *D. melanogaster* piRNAs are nearly 400 times more likely than random controls to be good matches. In addition to illustrating the different transposon silencing strategies employed by each organism, this result hints that the coverage of transposons achieved by *C. elegans* piRNAs might be comparable to coverage by random piRNAs. However, mismatch proportion is too simplistic a way of analyzing piRNA-target matches, since it does not take into account piRNA binding rules. A fair comparison to random controls demands a more refined analysis of *C. elegans* piRNA coverage.

### piRNA log-odds distance metric

How can sequence similarity be best measured in the context of piRNA binding? Previous studies have made an effort to systematize *C. elegans* piRNA targeting rules in order to locate piRNA binding sites. For example, the piRNA targeting site identification tool pirScan and the database piRTarBase utilize a targeting score which assigns differing penalties to mismatches or GU wobbles within or outside the seed region of the piRNA (17,18). While such a metric is useful for identifying targeting sites, the resulting score is highly dependent on the penalty values chosen and is difficult to interpret as a continuous parameter.

Instead, we developed a distance metric (see Methods) based on the log-odds probabilities of a match, mismatch, or GU wobble at each nucleotide position, obtained from the piRNA-target pairs experimentally confirmed by Zhang et al. (5). This metric is essentially the logarithm of a likelihood estimate of observing a given sequence of matches, mismatches or wobbles in a piRNA-target pair. When used to compare a piRNA and an mRNA, a smaller distance indicates a closer match, in a manner that reflects the importance of each nucleotide position to piRNA binding. The experimentally confirmed piRNA-target pairs used to construct the metric have a mean distance of 5.80 and a standard deviation of 2.22, which provides a useful benchmark to determine what distance constitutes a good match.

In order to verify that the distance metric is self-consistent and not overly dependent on the piRNA-target pairs used to construct it, we performed leave-one-out cross-validation, calculating the piRNA-target distances for each of those pairs while omitting them from the set used to calculate the probabilities incorporated into the distance (Supplementary Figure 1). We observed only small differences in the distance (0-3 units) when using these leave-one-out test sets instead of the full training set, indicating that the distance metric still gives similar results even when using a slightly different set of pairs as inputs.

### piRNA-transposon and piRNA-transcript targeting

The *C. elegans* piRNA system must silence transposons and unknown transgenes, while allowing self-genes to be expressed. If the expression of self-genes is protected by a separate licensing system, then the piRNA system could be largely agnostic to the identity of its targets. To what extent are piRNAs truly random with respect to the sequences they target?

To answer this question, we calculated the piRNA-target distance for all pairs of *C. elegans* piRNAs and their best-matching subsequence on each transposon, as well as the piRNA-target distance for all piRNAs and all best-matching subsequences of a random sample of 1,000 self-transcripts (Figure 3).

**Figure 3.**
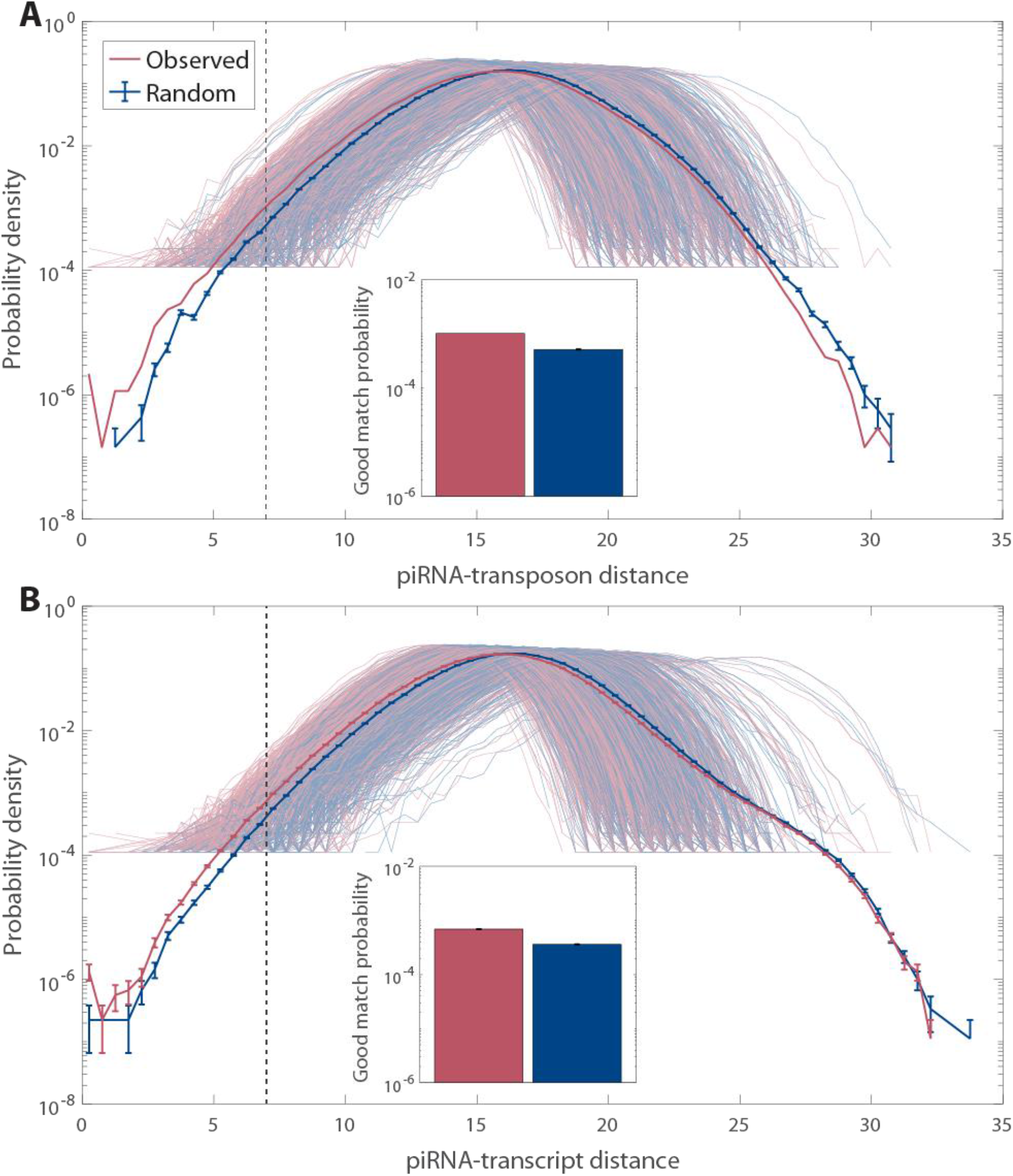
Bioinformatic analysis of *C. elegans* piRNA-transposon and piRNA-transcript pairing using the functional piRNA distance metric. **(A)** Probability distribution of the piRNA-transposon distance for every pair of *C. elegans* piRNAs (*n* = 17,849) and transposons (*n* = 752); results shown for real piRNAs and random control sequences. Pale curves indicate the distribution for each individual transposon; solid curve is the average over all pairs. (For piRNA-transposon distances that do not occur in the sample, no data is shown.) Inset: proportion of piRNA-transposon pairs where the best-matching site has a piRNA-mRNA distance of less than 7 (left of the dashed gray line in the main panel), for real piRNAs and random control sequences. Error bars indicate counting error. **(B)** Same as (A), but for a random sample of all *C. elegans* transcripts (*n* = 1,000) rather than transposons.

The probability distributions of distances appear very similar between real piRNAs and random controls, for both transposons and self-transcripts. We are particularly interested in the weight of the left (well-matching) tail of the distribution, to see whether real piRNAs are more likely than random controls to target a given class of mRNAs (Figure 3, inset). There are small differences in the proportion of “good” matches between the groups: transposons are slightly more likely to be targeted than self-transcripts, and real piRNAs are more likely than random controls to target both transposons and self-transcripts. However, none of these differences approach the amount of specific targeting observed in *D. melanogaster* (Figure 1), and are certainly insufficient to explain how transposons and self-transcripts are differentially silenced. We conclude that *C. elegans* piRNAs are functionally close to being random relative to their target sequences.

### Global targeting of arbitrary sequences

If *C. elegans* piRNAs are essentially random, then real piRNAs and random sequences should be able to target arbitrary nonself sequences with similar levels of reliability. Under this framework, the coverage of piRNAs depends on the number of distinct piRNA sequences: with enough random sequences, at least one is likely to be able to target a subsequence on any given mRNA. Is the number of *C. elegans* piRNAs (about 17,000) consistent with the required sequence coverage? To address this question, we calculated the mean piRNA-target distance for the best match between a set of random “piRNAs” and a random “gene,” as a function of the number of piRNAs and the length of the gene (see Methods). The average closest distance observed for the true number of *C. elegans “*piRNAs” (17,849) is similar to the distances of the experimentally confirmed piRNA-target pairs, suggesting that the number of piRNAs is tuned to allow sufficient coverage by what are effectively random piRNA sequences (Figure 4A).

**Figure 4.**
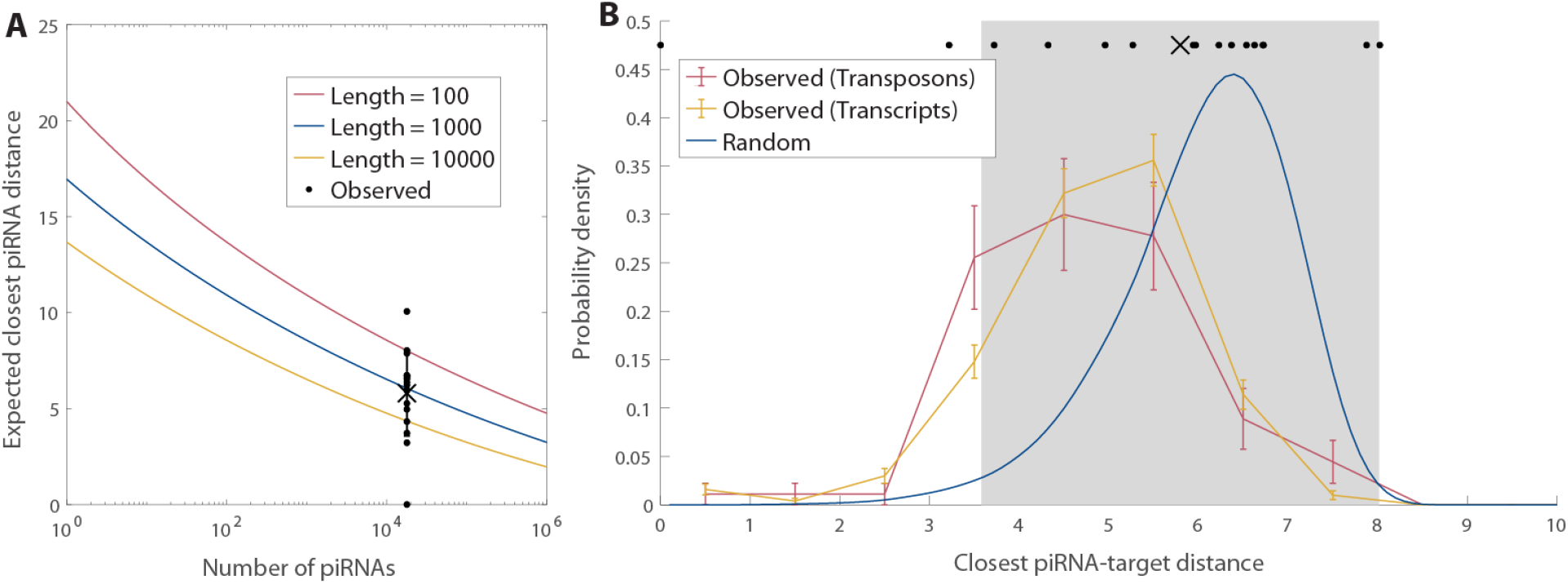
Comparison of actual targeting of genes by piRNAs to targeting of random “genes” by random “piRNAs.” **(A)** Predicted mean distance for the closest match between a “gene” (random sequence of length *L* = 100, 1,000, or 10,000), and any of the “piRNAs” (random 21 nt sequences) as a function of the number of distinct “piRNAs.” The data points plotted above the true number of *C. elegans* piRNAs (17,849) show the distances for the 17 actual piRNA-target pairs studied by Zhang et al. (5), with the cross and error bar indicating the mean and standard deviation. **(B)** Probability distribution of distance of the closest match for actual transposons and transcripts, of similar lengths, with the actual 17,849 *C. elegans* piRNAs. Red and yellow curves represent data for 800-1,200 nt *C. elegans* genes: transposons (*n* = 90, red) and a random sample of self-transcripts (*n* = 500, yellow). The blue curve shows the smoothed probability density of closest distances for a random “gene” of 1,000 nt and 17,849 random “piRNAs.” Error bars indicate counting error. The data points plotted above the probability distributions show the distances for the 17 actual piRNA-target pairs presented in (A), while the cross and gray shaded region show the mean and standard deviation, respectively.

We also wondered how the best-matching piRNA to a real transposon or self-transcript compared to what we would expect from random sequences. We restricted our attention to transposons and self-transcripts of approximately 1,000 nucleotides, and determined the data-based probability distribution of the distance between these sequences and their best-matching *C. elegans* piRNAs. We compared this distribution to that predicted by the fully random model, using the true number of *C. elegans* piRNAs and a 1,000-nucleotide target “gene” (Figure 4B). On average, the best-matching piRNA to a real transposon or transcript is a better match than the purely random model would predict (distances 4.8 ± 0.1 for self-transcripts, 4.7 ± 0.1 for transposons, and 6.1 for random sequences, with error indicating standard error of the mean). The difference between real and random piRNAs is much clearer than what we observed when considering all pairs of piRNAs and transposons (Figure 3). This is because focusing on the best-matching piRNA magnifies the small differences in the left tails of the distance distributions we found in Figure 3, which indicate how frequently piRNAs and mRNAs are good matches. However, the average best-matching piRNA-target distance for both real and random piRNAs is well within the range of typical distances observed for real piRNA-target matches — in fact, almost all of the density of both the experimental and random distributions is concentrated within this range. This result suggests that while piRNA targeting of real sequences is slightly better than random chance, random sequences would nonetheless be sufficient for targeting: the binding rules are tolerant enough that random “piRNAs” would be able to target a gene with the same accuracy observed in experimental piRNA-target pairs.

### piRNA-piRNA distance analysis

While *C. elegans* piRNAs appear nearly random with regards to the sequences they target, we also sought to determine how random they are with respect to each other. If piRNA sequences are self-avoiding, their coverage would be maximized with the minimum number of piRNAs. We therefore adapted the piRNA-target distance used earlier to measure piRNA-piRNA distance, where a smaller distance indicates more redundant coverage (see Methods). We calculated the distance between all pairs of *C. elegans* piRNAs, comparing the resulting probability distribution to that of random controls (Figure 5). The distributions appear similar, although the real piRNAs have a slightly larger left tail, corresponding to piRNAs with similar coverage of targets. We investigated this difference further by performing hierarchical clustering on both the real piRNAs and the random controls, grouping them into clusters with similar coverage (Figure 5, inset; see Methods). Large clusters are significantly more likely to occur in real piRNAs than in random controls, indicating that there are sets of piRNAs with very similar sequences.

**Figure 5.**
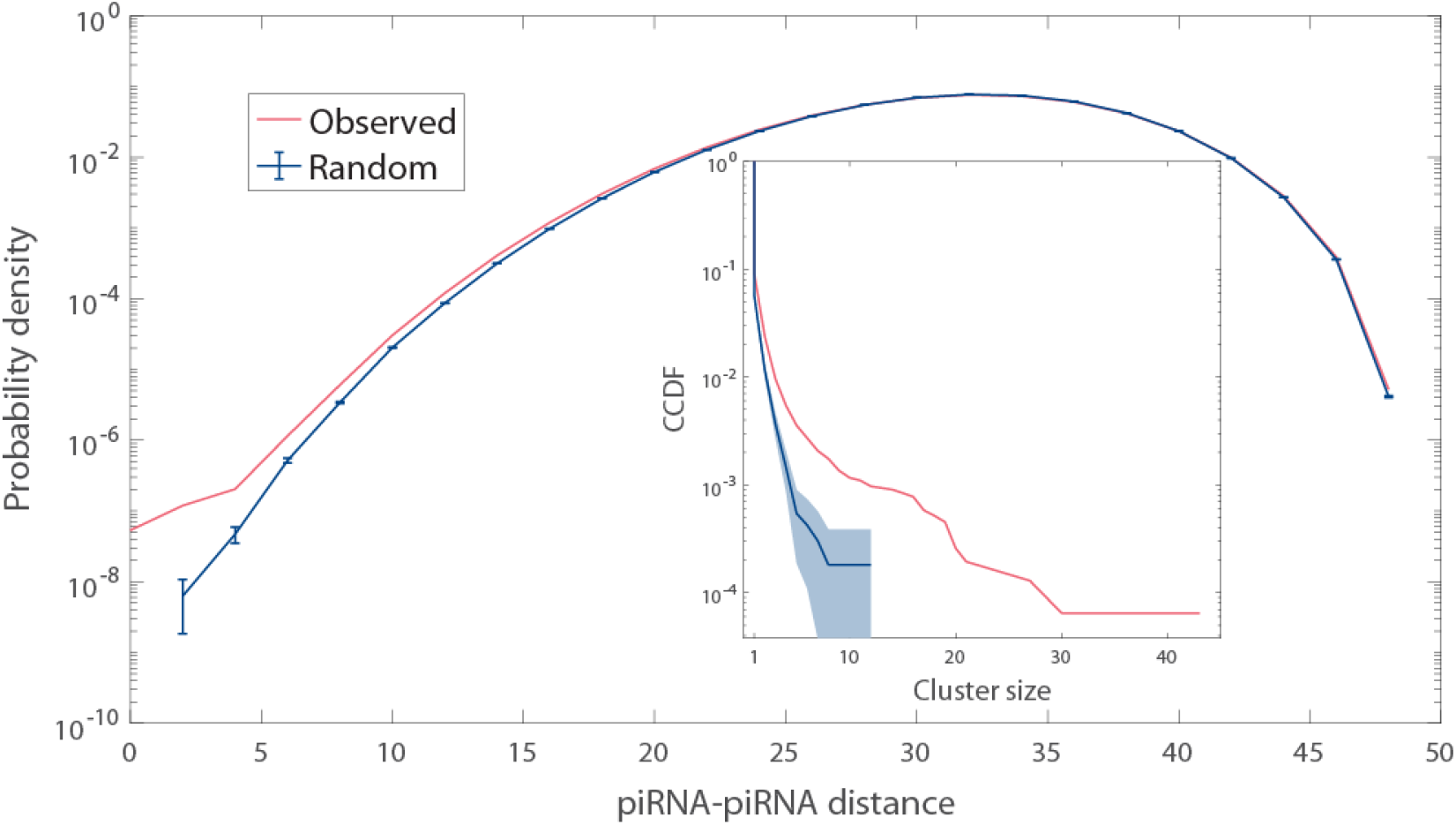
Sequence similarity among *C. elegans* piRNAs. Probability distribution of piRNA-piRNA distances; results shown for all 17,849 real piRNAs and for random control sequences. Error bars indicate counting error. Complementary CDF (CCDF) of cluster sizes for piRNAs and for random control sequences, as grouped via hierarchical clustering with single linkage (see Methods for details). The shaded region indicates a 95% confidence interval for the CCDF of random sequences.

Since real piRNA sequences are no more distant from each other than random sequences, they do not appear to be self-avoiding or “optimally” distributed. In fact, based on our clustering analysis, they are more likely to have *redundant* coverage than random sequences, which may be a result of new piRNA sequences being generated via duplication and modification of existing sequences. Altogether, *C. elegans* piRNA targeting does not appear functionally different from what would be achieved by the same number of random sequences.

## DISCUSSION

Using an original metric of piRNA-target distance, we compared piRNA targeting in *C. elegans* to that of random sequences. We found that piRNAs target both transposons and self-transcripts only slightly better than we would expect from fully random sequences; in fact, the same number of random sequences would be sufficient to reproduce the quality of targeting observed in experimental piRNA-target pairs. Furthermore, piRNA sequences are not self-avoiding or “optimally” distributed, and instead are slightly more redundant than random sequences. We conclude that *C. elegans* piRNAs achieve broad targeting by having a sufficient number of quasi-random sequences, which bind to targets with enough mismatch tolerance to cover all of sequence space.

Most of our analysis focused on comparing true *C. elegans* piRNAs and their targets to random sequences. The latter were generated by simply assigning global probabilities to each nucleotide — either equal to the nucleotide frequency in *C. elegans* piRNAs, or simply equal probabilities for each base. However, biological sequences are not random, possessing certain motifs or subsequences at higher or lower frequencies. Some of the differences we observed between random and real sequences can likely be ascribed to this fact, rather than to any specific targeting. For example, if a codon is more likely than others to appear in a sequence, a piRNA whose seed region targets this codon would raise the proportion of good matches relative to random controls without necessarily targeting a specific transposon. Such statistically enhanced targeting contrasts sharply with the precise targeting we noted in *D. melanogaster*, which utilizes piRNAs complementary to specific transposon sequences.

We analyzed targeting using an original piRNA distance metric (detailed in Methods). This metric is a log-odds score, corresponding to the logarithm of the likelihood estimate of observing a given sequence of matches, GU wobbles, and mismatches between a piRNA and its target. Based on experimental observations, the distance metric explicitly incorporates the assumption that canonical nucleotide matches at each position should pair at least as well as GU wobbles, and that GU wobbles in turn should be at least as favored as mismatches (5). The distance metric automatically incorporates the seed and supplementary regions via the position-specific data used to build the likelihood estimate. Thus the metric provides a continuous, data-based scale that takes into account the position-specific nuances of piRNA targeting. There are a variety of ways to define similar metrics, but the precise details of the distance used are unlikely to have affected our conclusions, as confirmed by our leave-one-out cross-validation testing.

Biologically, what is required for *C. elegans* piRNAs to achieve such broad coverage? We analyzed the number of distinct piRNA sequences as a key factor in determining coverage, suggesting that it has been evolutionarily tuned. One possible mechanism to change the number of sequences is duplication and modification of piRNAs, which may explain our identification of clusters of similar piRNA sequences. While our analysis took the piRNA binding rules as given, piRNA binding specificity is also biologically determined, likely by the PRG-1 Argonaute protein that combines with piRNAs to seek out target sequences. Studies of human Argonaute-2 (hAgo2) indicate that hAgo2 undergoes conformational changes as certain nucleotides of its guide miRNA bind, increasing its binding stability by exposing more guide RNA (19). Mutations in the PRG-1 Argonaute, then, could affect the number of matching nucleotides needed for stable binding to an mRNA and thereby the specificity of target identification. Since the number of piRNA sequences and the binding specificity are sufficient to determine coverage if the sequences themselves are arbitrary, investigating their coupling across similar systems in different species could shed light on how these systems function.

Because the *C. elegans* piRNA system is capable of targeting virtually all sequences, the organism must maintain a memory of self-sequences and protect them from silencing. This licensing system is not yet fully understood, but the mechanism of piRNA binding provides some clues: since a single transcript may have numerous, unpredictably spaced piRNA binding sites, physically blocking piRNA binding across the entire transcript would be difficult and likely unfeasible. An attractive alternative involves phase separation: licensed transcripts could be sequestered from the silencing machinery in physical space. Dodson and Kennedy posit that piRNA targeting occurs in perinuclear compartments of germ cells known as P granules, while amplification of the silencing response takes place in the nearby *Mutator* foci (20). P granules store newly transcribed mRNAs before releasing them into the cytoplasm for translation (21), and recent experiments show that during a silencing response, an enlarged P granule collects high amounts of the silenced transcript (22). The P granule has been found to contain both PRG-1 (8), which binds piRNAs, and the putative licensing Argonaute CSR-1 (23), which binds siRNAs complementary to most germline-expressed genes (10,11). These observations suggest a possible mechanism of licensing where newly transcribed mRNAs are allowed to bind to both PRG-1 and CSR-1 in the P granule, and whichever dominates determines whether transcripts are sequestered to the *Mutator* foci for downstream silencing or released from the P granule for translation (24). However, the precise mechanism of CSR-1 in licensing has not been confirmed, and other proposals for licensing systems exist: e.g. Zhang et al. argue that 10-base periodic A_n_/T_n_ sequence clusters act as licensing signals (5). Regardless of the details of the licensing mechanism, the ubiquity and random locations of piRNA targeting sites on transcripts means that physical separation is likely to play a key role in distinguishing silenced transcripts from those destined for translation.

Why might *C. elegans* have evolved a silencing system so broad that it necessitates a parallel licensing system? Across a variety of organisms, the evolution and counter-evolution of transposons and the piRNA system constitutes an “arms race” which drives rapid evolution and divergence between species (1). Indeed, *C. elegans* and its close relatives are the only nematodes to possess piRNAs; other nematodes employ different strategies of transposon silencing (25). One advantage of the *C. elegans* piRNA system over a “transposon library” system such as that of *D. melanogaster* is that it allows for rapid silencing of novel sequences, independent of sequence mutations in transposons or piRNAs. This idea is consistent with research suggesting that activation of transposons in *C. elegans* is associated with failures of the piRNA system, but not mutations in individual piRNA sequences (26). A rapid response may be especially important because *C. elegans* has a fixed number of cells, potentially making a takeover of the transcription machinery by a virus or transposon more costly to the organism. Another possible benefit is that a broad silencing system might enable *C. elegans* to dynamically control transcription, by changing the profile of what is being licensed (for example, by transcribing different guide siRNAs for CSR-1). In support of this hypothesis, germ cells in *C. elegans* which lack key P granule components undergo spurious differentiation pathways into other cell types, indicating a failure in transcriptional regulation (27). An understanding of the purpose of broad targeting by piRNAs in *C. elegans* could elucidate the role of piRNAs in other species. For example, mice possess two classes of piRNAs: pre-pachytene piRNAs with high complementarity to transposons, and pachytene piRNAs that possess much fewer clear matches to transposons in the genome yet still have regulatory roles (1,28). These pachytene piRNAs are responsible for broad purging of mRNAs during spermiogenesis (29), suggesting that multiple organisms employ the strategy of using a variety of arbitrary sequences to target a swath of mRNAs.

There are a variety of experiments that could further elucidate how *C. elegans* determines whether to silence or license transcripts, and measure the reliability and efficiency of this process. First of all, the piRNA-target distance implemented in this study could be further verified and refined by determining how effectively it predicts the probability of silencing. This can be done through a GFP reporter assay such as that used by Zhang et al., where a GFP transgene modified to lack piRNA targeting sites is introduced to the organism, along with a synthetic piRNA which partially matches a subsequence of the transgene (5). Changing the sequence of the synthetic piRNA relative to the modified GFP would allow silencing probability to be determined as a function of piRNA-target distance. Another key line of investigation is how *C. elegans* is able to rapidly classify transcripts as self or nonself: transcripts destined for expression only spend about 15 minutes in the P granule before being released to the cytoplasm, suggesting that Argonaute binding and target identification must happen in this timeframe (21). A promising line of research to address this question is analyzing the mRNA scanning properties of the Argonautes involved in this system, PRG-1 and CSR-1, to estimate their search speed — which would also determine whether different Argonautes have been evolutionarily tuned for different levels of mismatch tolerance. The FRET assay used by Chandradoss et al. to estimate the dwell time of the hAgo2-miRNA complex on a target subsequence with a given number of matches would be a useful tool to address this question (30). These experiments and others would help lead us toward a deeper quantitative understanding of how *C. elegans* utilizes small RNAs to efficiently differentiate self and nonself transcripts.

## AVAILABILITY

The resources used in this study are freely accessible online, detailed in the Computational Resources section of Materials and Methods.

## ACCESSION NUMBERS

We have no accession numbers to report.

## ACKNOWLEDGEMENT

We thank our colleagues at Princeton University for lending their expertise while we developed this study: Coleen Murphy and Rachel Kaletsky for their guidance on *C. elegans* and piRNAs, Joshua Riback for our discussion about phase separation, and Cameron Myhrvold for our discussion on experimental imaging methods. We also thank Wei-Sheng Wu at National Cheng Kung University for providing assistance with the pirScan tool he helped develop when we were performing initial investigations into this system.

