## Supplementary Figure 1 for "piRNAs of *Caenorhabditis elegans* broadly silence nonself sequences through functionally random targeting"

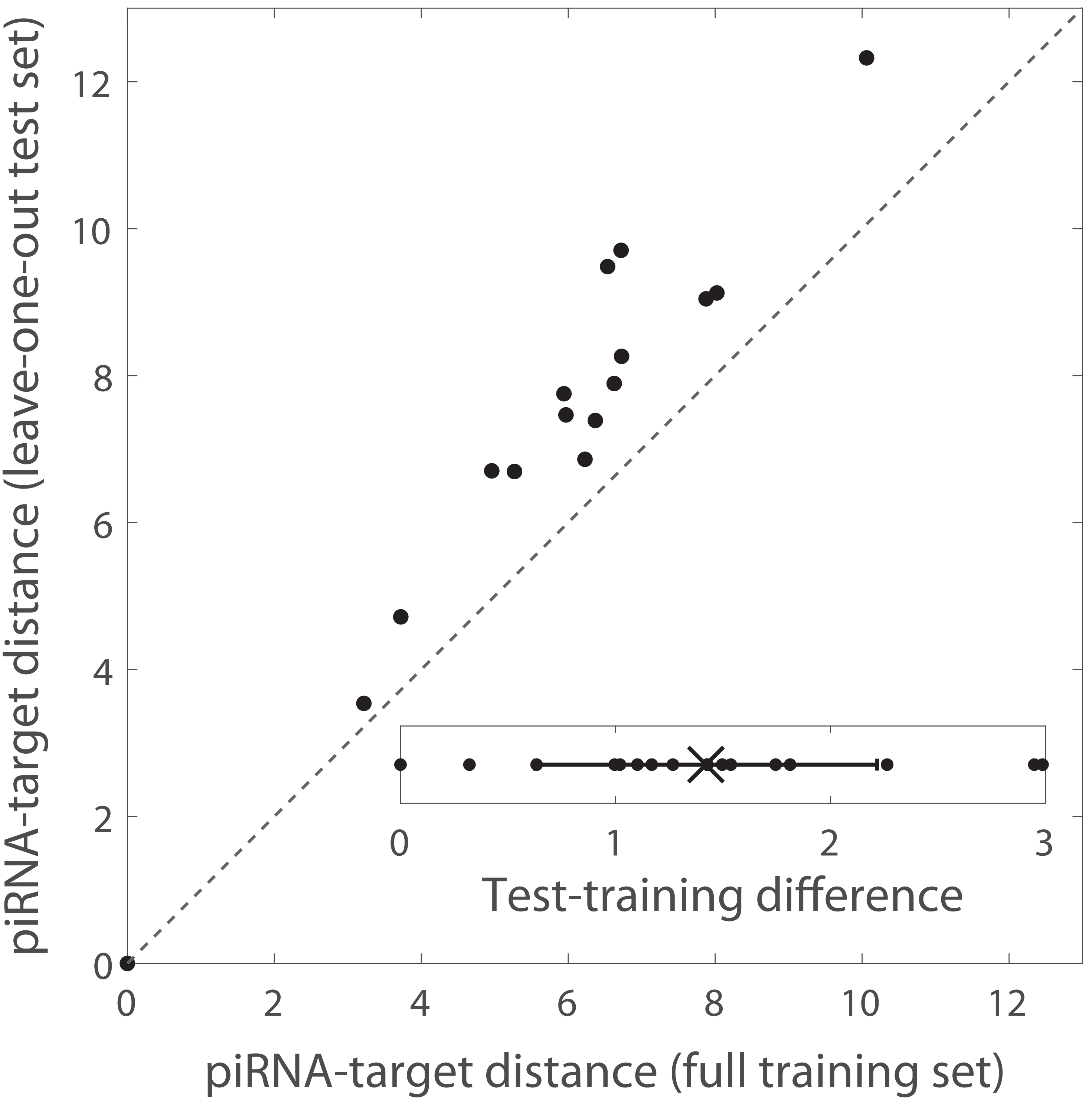


Supplementary Figure 1. Leave-one-out cross-validation of distance metric. Comparison between piRNA-target distances of known piRNA-target pairs using the full training set (the piRNA distance utilized in the rest of the paper, incorporating all known pairs) or leave-one-out test sets (where the pair being measured is omitted when calculating the log-odds probabilities). The dashed line indicates perfect equivalence. Inset: the difference between the distance calculated between each piRNA-target pair in the leave-one-out test set and in the full training set, for each of the 17 experimentally confirmed piRNA-target pairs. The cross and error bar indicate mean and standard deviation.
